# The importance of m^6^A topology in chicken embryo mRNA; a precise mapping of m^6^A at the conserved chicken β-actin zipcode

**DOI:** 10.1101/2022.03.04.483006

**Authors:** Francis Baron, Mi Zhang, Nathan Archer, Eleanor Bellows, Helen M. Knight, Simon Welham, Catrin S Rutland, Nigel P. Mongan, Christopher J. Hayes, Rupert G. Fray, Zsuzsa Bodi

**Author notes:** Corespondance: Rupert G. Fray,; Christopher J. Hayes,; Zsuzsa Bodi.

## Abstract

*N6*-Methyladenosine (m^6^A) in mRNA regulates almost every stage in the mRNA life cycle, and the development of methodologies for the high throughput detection of methylated sites in mRNA using m^6^A-specific methylated RNA immunoprecipitation with next-generation sequencing (MeRIPSeq) or m^6^A individual-nucleotide-resolution cross-linking and immunoprecipitation (miCLIP) have revolutionized the m^6^A research field. Both of these methods are based on immunoprecipitation of fragmented mRNA. However, it is well documented that antibodies often have nonspecific activities, thus verification of identified m^6^A sites using an antibody-independent method would be highly desirable. We mapped and quantified the m^6^A site in the chicken β-actin zipcode based on the data from chicken embryo MeRIPSeq results and our RNA-Epimodification Detection and Base-Recognition (RedBaron) antibody independent assay. We also demonstrated that methylation of this site in the β-actin zipcode enhances ZBP1 binding *in vitro*, whilst methylation of a nearby adenosine abolishes binding. This suggests that m^6^A may play a role in regulating localised translation of β-actin mRNA, and the ability of m^6^A to enhance or inhibit a reader protein’s RNA binding highlights the importance of m^6^A detection at nucleotide resolution.

## Introduction

Amongst more than 100 modified RNA nucleotides, *N6*-Methyladenosine (m^6^A) is the most abundant internal modification in eukaryotic mRNA. m^6^A regulates almost every stage of the mRNA life cycle, with important regulatory roles in splicing (Xiao et al. 2016), polyadenylation (Ke et al. 2015), nuclear export (Roundtree et al. 2017, Lesbirel et al. 2018), stability (Wang et al. 2014), translation (Wang et al. 2015, Zhou et al. 2015, Meyer et al. 2015) and degradation (Wang et al. 2014, Du et al. 2016). m^6^A is essential for normal development of eukaryotic organisms (Zhong et al. 2008, Geula et al. 2015), and abnormal levels of m^6^A have been associated with diseases including various types of cancer (Zhang et al. 2016, Lu, et al. 2017, Chen et al. 2018).. Transcriptome wide, between 0.2 and 0.4 % of adenosines are m^6^A modified (Dominissini et al. 2012, Meyer et al. 2012, Schwartz et al. 2014), depending on tissue or cell type. However, the modification is unevenly distributed in mRNA transcripts and is predominantly localised in the 3′ UTR near the stop codon (Dominissini et al. 2012, Meyer et al. 2012), usually within the consensus sequence motif DRACH (D=G/A/U, R=G/A, H=A/U/C).

There are a number of methods for the transcriptome-wide detection of m^6^A. The most commonly used methods are m^6^A Seq/MeRIP-Seq (Dominissini et al. 2012, Meyer et al. 2012, Schwartz et al. 2014) and miCLIP (Linder et al. 2015), however, these methods are unable to unambiguously distinguish m^6^A at specific nucleotide sites or to quantify the proportion of a particular gene’s transcripts which contain the modification at a specific site. Both methods require the use of an anti-m^6^A antibody, and these antibodies can exhibit off target activities, commonly targeting purine rich, non-methylated regions of the RNA, potentially resulting in false positives (Helm et al. 2019). Alternative methods of m^6^A mapping have been tested, including the use of reverse transcriptases (Harcourt et al. 2013, Wang et al. 2016, Aschenbrenner et al. 2018) and modified nucleotide triphosphates (Hong et al. 2018), as well as the use of inhibition of endoribonuclease MazF to cut RNA at ACA sites when methylated (Imanishi, et al. 2017, Zhang et al. 2019). None of the above methods, gives a representative picture of the whole methylome with high certainity. Third generation sequencing technologies such as Oxford Nanopore and single molecule real time (SMRT) sequencing are rapidly improving and show great promise (Liu et al. 2019, Vilfan et al. 2013) for m^6^A detection, however, these methods are also limited in their accuracy due to the lack of good synthetic training sets reflecting the biological diversity of m^6^A-contexts *in vivo* and antibody independent verification methods.

There is a recognized need for a sensitive biochemical method for transcript-specific m^6^A detection and quantification. To date, there are only two methods capable of this, SCARLET and SELECT (Liu et al. 2013, Xiao et al. 2018). However, SCARLET is technically difficult, time consuming, and requires large quantities of input RNA. For these reasons, the SCARLET method is not routinely used. SELECT is claimed to be a very simple, low input, qPCR based method. However its accuracy is dependent on the very precise quantification of the input RNA concentrations. SELECT can be quantitative in determining m^6^A/A ratios at a specific site. However, for this purpose, a precise quantification of the target transcript in the input must be performed alongside a calibration curve for the m^6^A/A fractions in the sequence context of the assumed m^6^A position. These additional steps make SELECT more laborious than a ‘one tube’ experiment, and potentially reduce accuracy.

Actin is one of the most conserved proteins within metazoans and its transcript is m^6^A methylated in human and mouse (Dominissini et al. 2012). Amongst the different isoforms, β-actin is a cytoplasmic actin that is highly regulated both spatially and temporally and plays an essential role during development. It is also involved in cell shape changes, protein trafficking, cell division, chromatin remodeling and regulation of transcription (Vedula and Kashin 2018, Lehtimaki et al. 2017, Luxenburg and Geiger, 2017, Viita and Vartiainen, 2017, Almuzzaini, et al. 2016). β-actin mRNA has been shown to localize to the leading edge of chicken embryo fibroblasts and to the extending neuronal growth cones (Lawrence and Singer 1986, Zhang et al. 1999). The spatial targeting of β-actin mRNA is under the control of the zipcode sequence, located in the 3′UTR of the transcript. The zipcode sequence is responsible for recruiting the highly conserved KH (hnRNP K homology) domain zipcode binding protein, ZBP1. The 28 nucleotide (nt) zipcode contains the highly conserved GGACU sequence, and this motif is essential for the KH domain binding (Chao et al. 2010, Nicastro et al. 2017). This same sequence happens to be the canonical consensus sequence for m^6^A mRNA methylation in most eukaryotes, and is methylated in mouse and human (Dominissini et al. 2012, Liu et al. 2013). The conserved zipcode sequence between eukaryotes and the site of m^6^A at this position suggests a link between actin localization and mRNA methylation.

Here we demonstrate that the chicken *ACTB* zipcode sequence has m^6^A sites which we accurately map and quantify using a low input, quantitative **<u>R</u>**NA-**<u>E</u>**pimodification **<u>D</u>**etection and **<u>Ba</u>**se-**<u>R</u>**ecogniti**<u>on</u>** ‘RedBaron’ verification method. We also demonstrate that, the presence of m^6^A can either enhance or abolish ZBP1 binding *in vitro* depending on its precise site within the zipcode sequence.

## Results

### Transcriptome wide detection of m^6^A in *Gallus gallus*

The two core components of the m^6^A methylase writer complex, *METTL3* and *METTL14*, are 78.23% and 93.94% identical, respectively between chicken and mouse (Supplementary Data S1). Therefore, we were interested in testing how much m^6^A is present in the chicken transcriptome, and how it is distributed across the transcriptome compared to other vertebrates. Initially we measured the global levels of m^6^A in chicken poly(A) RNA from embryos and chicken embryonal fibroblast cells using the two-dimensional thin layer chromatography (TLC) method (Zhong et al. 2008). The m^6^A to A ratios (following a G) (Figure 1A) were very similar to that of published values in mouse and human (Liu et al. 2020).

**Figure 1.**
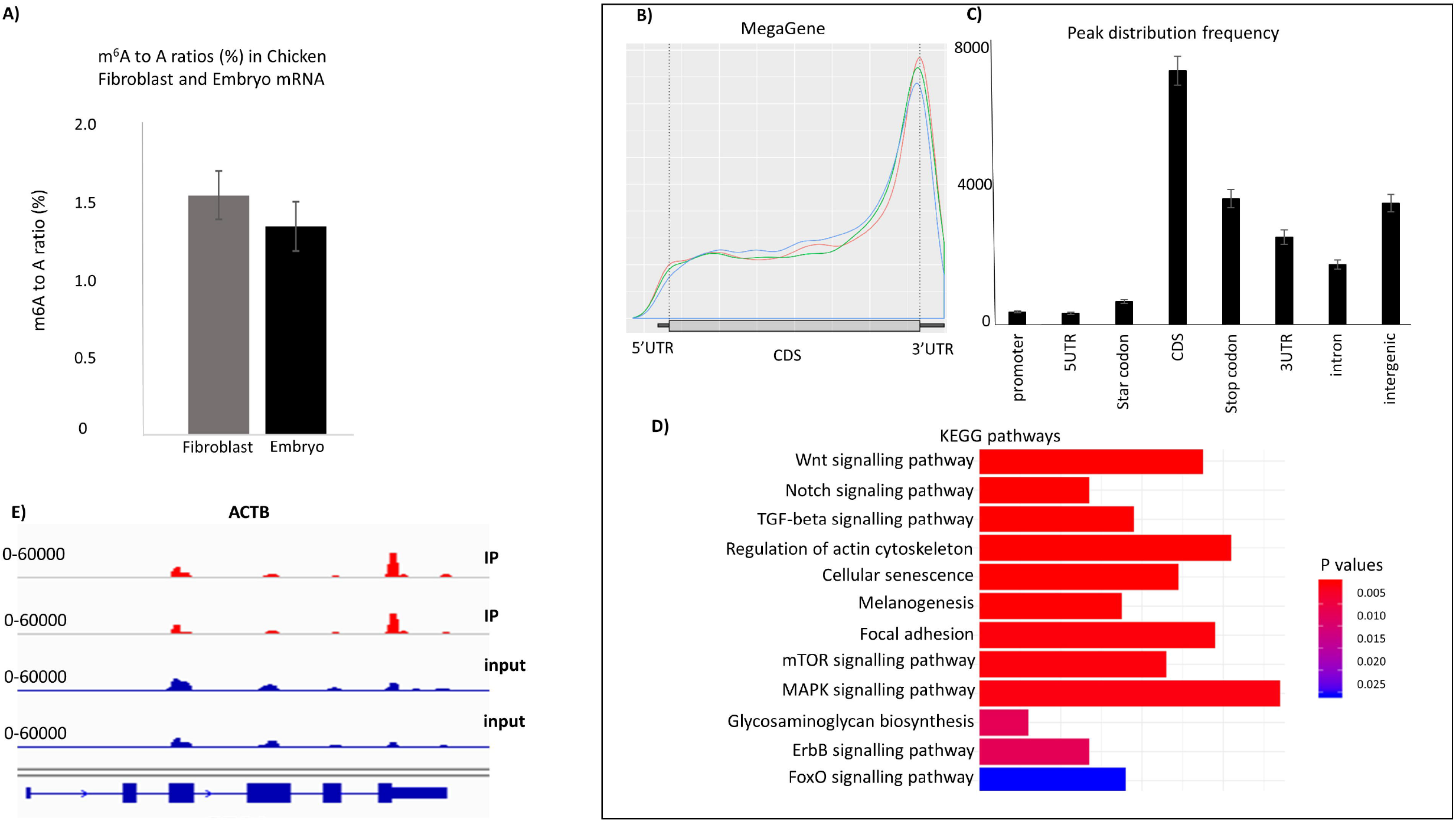
The topology of m^6^A in the chicken transcriptome. A) m^6^A levels in chicken embryo (HH27) and chicken embryonal fibroblast; for each experiment 3 biological replicates were used, the difference between the fibroblast and embryo is not significant (T-test, p = 0.173, one tailed).B) mRNA metagene plot from the RNAmod analysis of the MeRIPSeq data from 3 biological replicates of chicken embryo (HH27). The Y-axis represents the density distribution of coverages. C) Modification sites/peak distribution on different gene features. Y-axis represents the frequency of peaks/sites (number of peaks/sites) while x-axis represents different gene features. The error bars represent the standard deviations for 3 biological replicates. D) KEEG pathway enrichment for 3 replicates shows the top 12 most enriched pathways (data are created using the RNAmod platform http://61.147.117.195/RNAmod). The colour scale represents the enrichment p-value. E) Integrative Genomics Viewer (IGV) tracks of MeRIP-seq, upper panel and RNA-seq, lower panel read distribution of ACTB mRNA.

To determine the topology of m^6^A sites at transcriptome level we carried out a MeRIPSeq experiment on poly(A) fractions isolated from chicken embryos. m^6^A peaks were identified and functionally annotated using the RNAmod portal (http://61.147.117.195/RNAmod/) (Liu and Gregory, 2019). Only m^6^A peaks with cut off values for significance p ≤ 0.05, and four-fold increase in IP vs input were used (sample 1, 16989 peaks; sample 2, 13331 peaks; and sample 3, 9182 peaks). In the final gene matrix 4332 peaks were identified which were represented at least in two replicates (Supplementary Figure S1). The topology of the m^6^A deposition in the chicken transcriptome represented by the metagene analysis is very similar to mouse and human (Dominissini et al. 2012, Meyer et al. 2012). This analysis showed that most m^6^A peaks are concentrated around the 3′end of transcripts (Figure 1B) The peak distribution frequency in the 5′ UTRs is 10 fold lower compared to those in CDS and in the stop codon-3′UTR regions (Figure 1C). The gene type statistics showed that most of the peaks are found in protein coding transcripts (Supplementary Figure S2).

A pathway enrichment analysis using all significant methylation peaks with four-fold or greater increase indicated that several KEGG pathways characteristic for chicken stage HH27 (Hamburger and Hamilton 1951) were enriched in the m^6^A methylated transcript population (Figure 1D). One of the most significantly enriched pathways identified was the ‘Regulation of actin cytoskeleton’. 22 methylated transcripts, including *ACTB* belong to this pathway (Supplementary Table S1). Furthermore, using a conserved set of transcripts between mouse and chicken, and only those transcripts that were methylated at 3′ends in both species, we identified both, *ACTB* (chicken) and *Actb* (mouse) homologs (Supplementary Data S2) The KEGG pathway enrichment for the chicken 3′ UTR methylated conserved transcripts also identified the ‘Regulation of actin cytoskeleton’ as one of the top enriched pathways (Supplementary Table S2). The methylation peaks in chicken *ACTB* map within the zipcode binding sequence, immediately after the stop codon in the 3′UTR (Figure 1E). The GGACU site in the β-actin zipcode was previously found to be methylated in mouse and human (Dominissini, et al. 2012, Liu et al. 2013). However, m^6^A peak summits from our three experimental repeats did not align exactly over the GGACU sequence in the β-actin 3′UTR region thus demonstrating the limitations of MeRIP data, which were unable to pinpoint the precise position of m^6^A in the zipcode sequence. Knowing the precise position of m^6^A is important, as the zipcode sequence contains several **<u>A</u>**s that could be targets for methylation, and methylation at different sites might influence ZBP binding, and thus effect transcript fate, in different ways.

### Effect of zipcode methylation on ZBP1KH3-KH4 binding

The presence of the zipcode sequence is necessary for the β-actin mRNA subcellular localisation. β-actin is both structurally and functionally highly conserved between vertebrate species. Moreover, there is a conservation of the presence of m^6^A in the zipcode sequence of mouse, human (Dominissini et al. 2012, Meyer et al. 2012, Liu et al. 2013) and in chicken embryo β-actin transcripts (Figure 1E). Thus, the conservation in the m^6^A topology at the zipcode sequence suggests this modification may be functionally important for the spatial expression of β-actin, facilitated by ZBP1 binding.

The presence of m^6^A in the zipcode was verified for human β-actin mRNA using the SCARLET method. The precise position of m^6^A modification was the central adenosine of the ‘GG**<u>A</u>**CU’ sequence motif (position 1216, HeLa β-actin mRNA) and 21% of As in this position were m^6^A (Liu et al. 2013). This motif is an essential sequence within the 28 nt zipcode (Figure 2A) for binding and stabilizing of KH4, one of the four KH domains in the chicken ZBP1 protein, while an ACACCCC motif downstream to the GGACU is essential for KH3 binding (Chao et al. 2010, Nicastro et al. 2017) (Figure 2A, B). We hypothesized that methylation of the β-actin zipcode plays an important role in recruiting the ZBP1 protein. As the core ZBP1 binding chicken zipcode sequence contains several potential m^6^A sites, and it is not possible to unambiguously determine the precise position of m^6^A modifications from MeRIPSeq results alone (Helm et al. 2019) we wanted to test the effect of m^6^A presence at several AC sites for their influence on ZBP1 binding.

**Figure 2.**
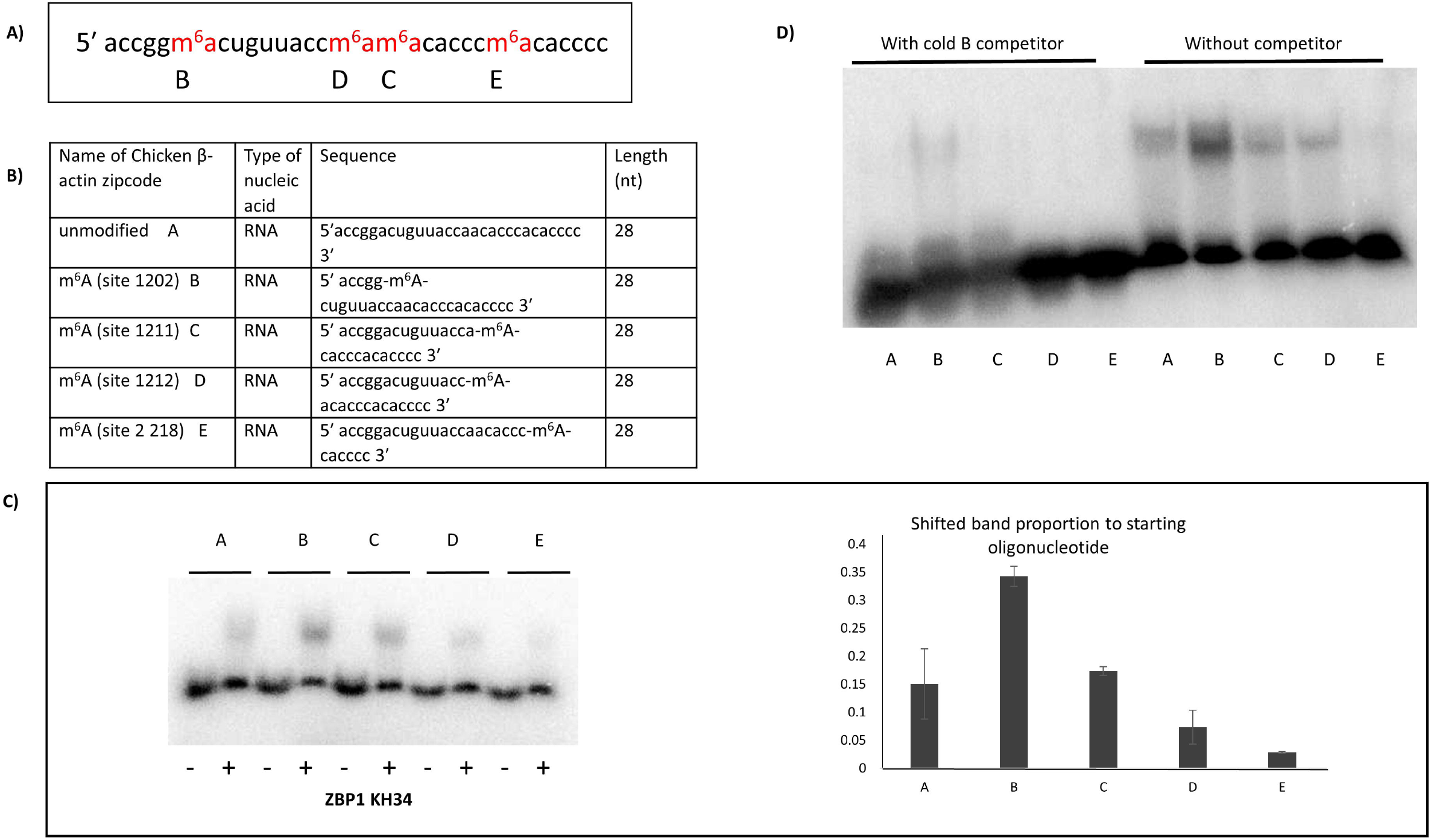
Methylation in the β-actin zipcode changes ZBP1 binding. A-B) Positions and labelling of the assayed m^6^A sites in the chicken β-actin core zipcode sequence. C) Binding assay of the ZBP1KH34 domain to the zipcode oligonucleotide containing m^6^A at the different positions. A, B, C, D, E refers to the m^6^A positions presented in (A) and (B) sub figures. The bar chart shows the quantification of the shifted RNP complexes. The error bars represent the standard error from 3 replicates. B and E oligomers are significantly different from A oligomer (T-test for A-B, p = 0.036; A-E, p = 0.044; B - E, p = 0.026; one tailed). D) Competition assay using cold B oligonucleotide.

Thus, in the first instance we synthetized a series of the core 28 nucleotide zipcode RNA sequences in which we replaced candidate A sites with m^6^A (within nucleotides 1-28) (Figure 2A, B). To test how m^6^A in different positions influences the ZBP1 binding, we performed gel shift assays using synthetic zipcode RNA oligonucleotides as previously described (Chao et al. 2010). The truncated ZBP1protein containing only KH3 and KH4 domains maintains the binding properties of the full length protein (Chao et al. 2010). Using this truncated KH3-KH4 ZBP1 in combination with different methylated versions of the zipcode sequence (Figure 2A), we showed that replacing A with m^6^A in the GG**<u>A</u>**CU motif resulted in a stronger ZBP1 binding (Figure 2C, D). This result is supported by similar observation from Huang et al., showing that IGF2BP1, the human homologue of ZBP1, is an m^6^A reader protein and stabilizes m^6^A harbouring transcripts (Huang et al. 2018), although these authors ere not looking at zipcode-specific binding and did not test the influence of m^6^A within a zipcode sequence context. All other m^6^A modifications in our synthetic zipcode oligonucleotides were neutral or negative in their effect on ZBP1 binding. When m^6^A replaced the A at position 22 in the **<u>A</u>**CACCCC (E oligonucleotide) sequence motif (Figure 2C), the methylation almost completely abolished the ZBP1 binding to the core zipcode sequence. Both motifs were previously reported to be essential for ZBP1 binding (Chao et al. 2010). A cold competitor that had an m^6^A in position 6 (GG**<u>m^6^A</u>**CU) out-competed all methylated and non-methylated zipcode sequences in the RNA-protein complexes (Figure 2D), further demonstrating that m^6^A in the GGACU context has the strongest binding to ZBP1.

### The RedBaron method for site specific detection and quantification of m^6^A

We developed the RedBaron method as SCARLET is technically difficult, time consuming, and requires large quantities of input RNA. Likewise, the SELECT method requires several qPCR steps for determining target transcript concentrations and creating calibration curves for m^6^A to A ratios in the sequence context where the m^6^A mark is to be assayed. RedBaron has only 3 simple steps. First, a chimeric oligonucleotide directs RNase H cleavage, which is similar to SCARLET. However, unlike SCARLET, the site-specific hydrolysis of the phosphodiester bond is designed to occur immediately 3′ to the A/m^6^A candidate site, leaving a 3′ OH. Second, a 5′ ^32^P radiolabeled DNA oligonucleotide is splint ligated using the 3′OH of the A/m^6^A candidate nucleotide, forming a chimeric RNA-DNA oligonucleotide. Third, the chimeric nucleic acid is digested into 3′ nucleoside monophosphates and two-dimensional thin layer chromatography (2D-TLC) is used to quantify the relative levels of adenosine and m^6^A (Figure 3). This method also avoids gel purification, dephosphorylation and labelling of all exposed RNA 5′ ends (which are required steps for SCARLET).

**Figure 3.**
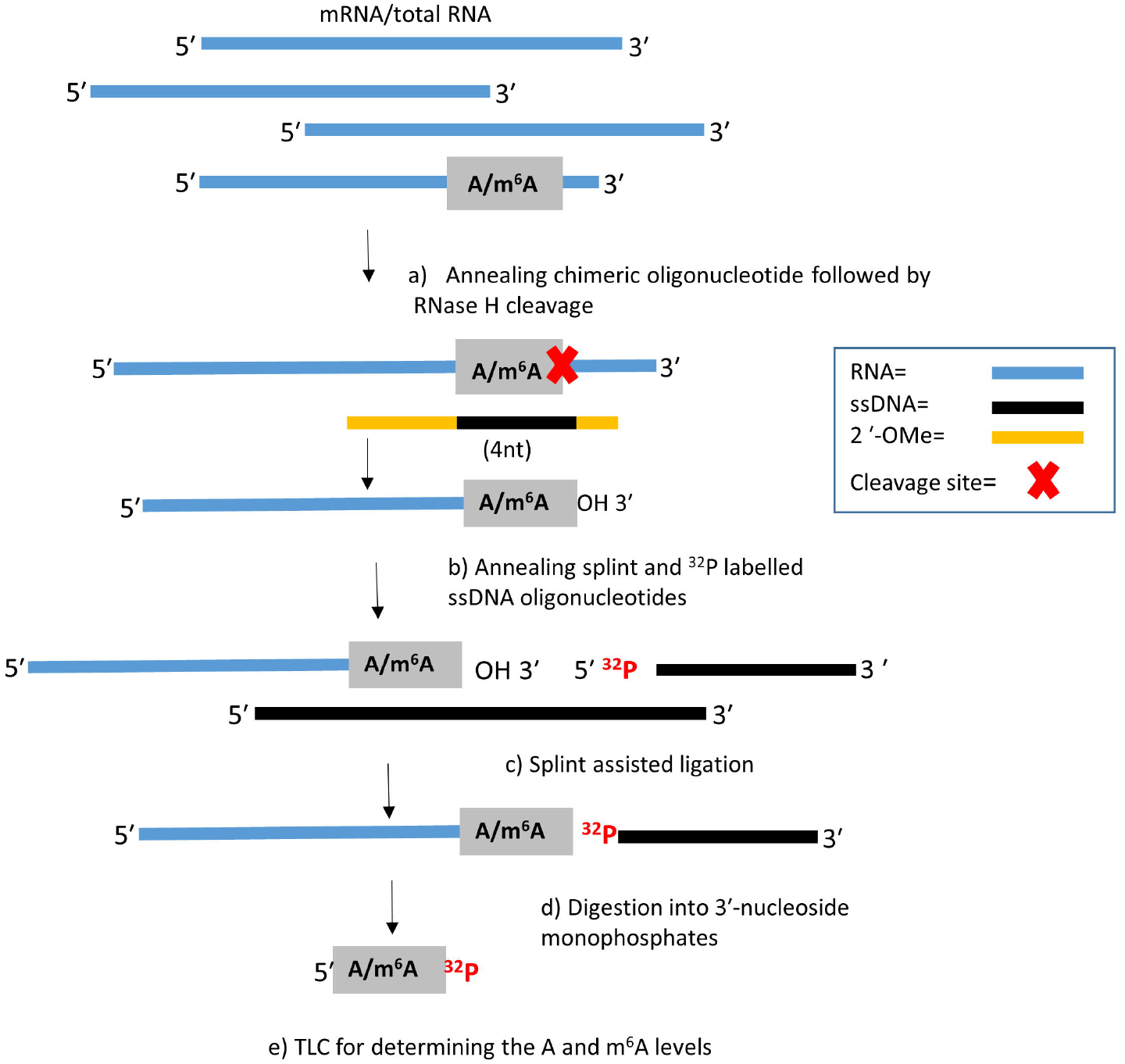
The RedBaron method of m^6^A detection: A chimeric oligonucleotide was used to target an RNase H cleavage of phosphodiester bond immediately 3′ to the A/m^6^A candidate site (A and B). A ^32^P radiolabeled single stranded DNA oligonucleotide is ligated to the 3′ of the A/m^6^A candidate nucleotide (C). The RNA is digested into 3′ nucleoside monophosphates (D). The relative levels of m^6^A to adenosine are quantified by 2D-TLC (E).

To demonstrate that the RedBaron method is able to accurately detect m^6^A, we synthesised two oligonucleotides containing either A or m^6^A at a specific position (Figure 4A). In the first instance we applied the RedBaron protocol to the synthetic m^6^A and A RNA oligonucleotides in two separate experiments. The 3′ nucleoside monophosphates (Ap and m^6^Ap) generated using the RedBaron method run slightly further in both the first and second dimension than the 5′ nucleoside monophosphates. For this reason, prior to the detection of radiolabeled nucleotides using TLC method, we spiked in an equimolar mixture of pA, pG, pC, pU nucleotides to determine the correct orientation of 3′ adenosine monophosphate and 3′ *N*6-methyladenosine monophosphate (Figure 4B). This allows easy distinction between A and m^6^A and the spiked in 5′ nucleotides (Figure 4B). Next, we tested the accuracy of the method for quantifying m^6^A amounts in a mixture of synthetic m^6^A modified and unmodified oligonucleotides. Using varying ratios of the unmodified and m^6^A modified oligonucleotides we demonstrated that the RedBaron method is able to accurately measure levels of m^6^A across a wide range of input values (Figure 4C). Thus, this method is quantitative and site specific in a synthetic system.

**Figure 4.**
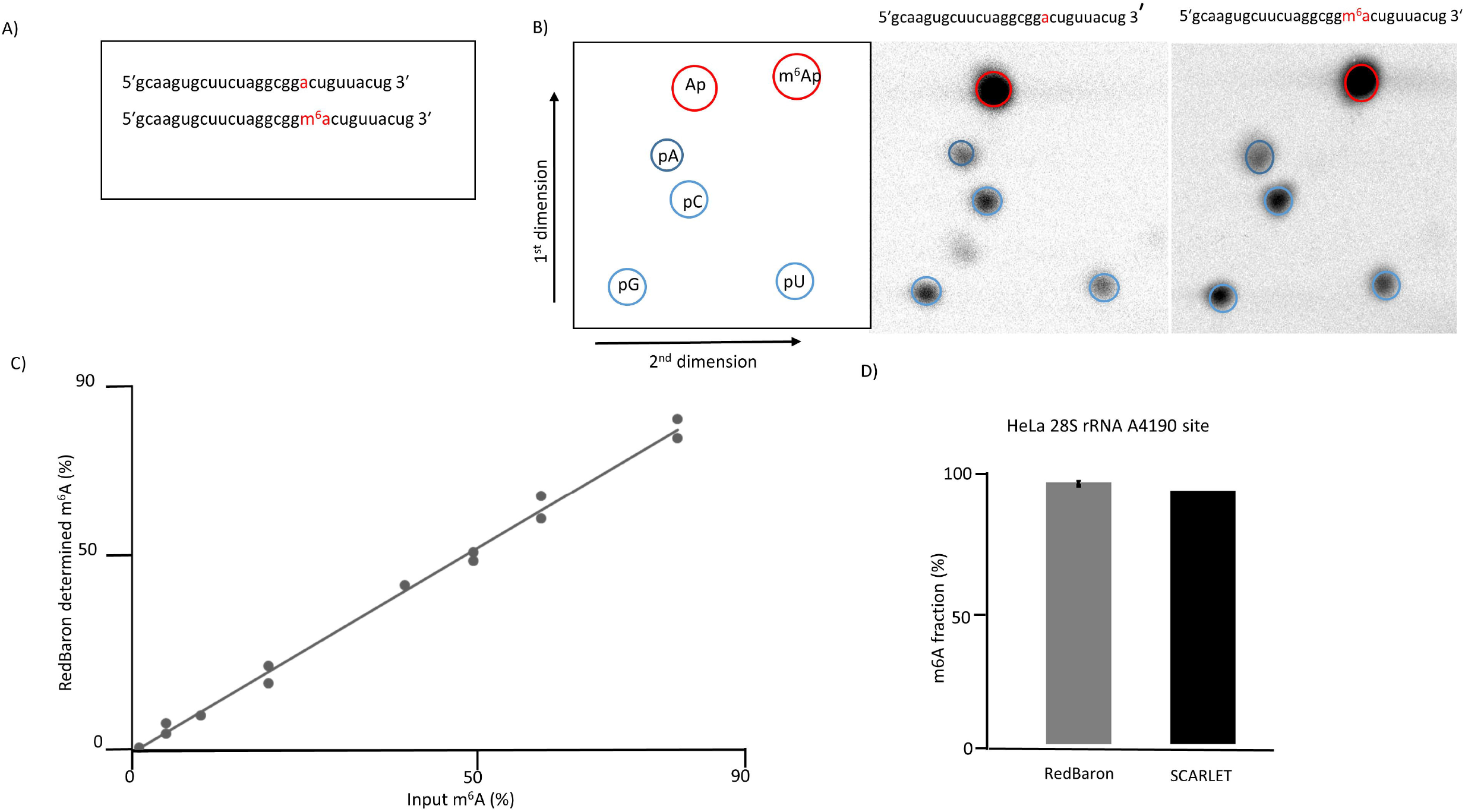
Detection and quantification specific sites using the RedBaron method. A) Synthetic RNA oligonucleotides containing either A or m^6^A at position 19. B) Two-dimensional TLC analysis is used to differentiate between the AP and m^6^AP nucleoside monophosphates (red) generated by the RedBaron method. Left: schematic picture of the TLC plate; Center: unmodified RNA template; Right: m^6^A modified RNA template. The 5′ nucleoside monophosphates (pA, pG, pC, and pU) are used as reference molecules (blue). C) The m^6^A/A ratios from the TLCs are accurately representing the concentration ratios of the two synthetic oligonucleotide mixes. D) The RedBaron analysis of HeLa 28S rRNA site A4190 showed an average of 99% m^6^A, observed from 3 repeats (99.25 %, 99.21%, 99.28%). This is similar to the SCARLET method’s 96% reported in Liu *et* al. 2013. Error bars for the RedBaron method show the standard deviation from three replicates.

Next, we chose the Human 28S rRNA for testing the RedBaron method’s sensitivity and accuracy within native RNA molecules. Human 28S rRNA contains only one significant m^6^A modification. Using the SCARLET method, Liu et al. (2013) observed 96% m^6^A at this site in HeLa RNA. Consistent with this result, we observed 99 % m^6^A at this site in HeLa RNA using the RedBaron method (Figure 4D). Over the three experimental repeats, we observed almost no variation in m^6^A levels. Thus, we conclude that the RedBaron method is quantitative and accurate using *ex vivo* samples.

### Site-specific detection and quantification of m^6^A in β-actin mRNA

Current MeRIPSeq or miCLIP methods are unable to precisely and unambiguously identify specific m^6^A locations at nucleotide resolution, due to the intrinsic limitations of the anti-m^6^A antibody specificity on which such methods depend (Helm et al. 2019). This is particularly true for transcripts with low level m^6^A. Since we observed that ZBP1 binding is dependent on the topology of m^6^A in the zipcode oligonucleotides *in vitro*, we wanted to test the presence of m^6^A at different positions *in vivo* and *ex vivo*. Therefore, we applied the RedBaron method for site specific identification and quantification of m^6^A sites in the chicken β-actin zipcode region. We used poly(A) enriched mRNA from chicken embryos and from chicken embryonal fibroblast cells. In the first instance we tested the methylation status of the A in the GG**<u>A</u>**CU context (Supplementary Figure S3A). We found 13% m^6^A at this site in chicken embryos, whilst chicken fibroblasts gave a higher value of 26.5% (Supplementary Figure S3B). As the global methylation levels are very similar between chicken embryo and fibroblast cells (Figure 1A) the difference in m^6^A levels at the zipcode sequences suggest a functional importance. When we tested the **<u>A</u>**CACCCC (position 22, E oligonucleotide) position using the RedBaron method we did not find any detectable m^6^A at this site (Supplementary Figure S3B). Thus, we can conclude that this ZBP1-suppressive position is not methylated *in vivo* at the developmental stage tested in chicken embryo.

## Discussion

### The importance of m^6^A position for RNA binding proteins

MeRIPSeq data from chicken embryo revealed that the transcript of one of the most conserved genes among vertebrates, β actin, was methylated near to the stop codon over the zipcode sequence, as is also seen in mammals. As the zipcode region determines the subcellular localization of actin mRNA via binding to ZBP1 (Chao et al. 2010) we tested how the presence of m^6^A influenced the binding properties of the ZBP1 KH3-KH4 domains that have previously been shown to be responsible for zipcode recognition (Chao et al. 2010). We used a series of synthetic oligonucleotides harbouring m^6^A at different sites in the core 28 nucleotide conserved chicken zipcode sequence. These experiments revealed that the presence of m^6^A in the zipcode GG**<u>A</u>**CU sequence enhanced ZBP1 binding. Our results are supported by the study on the human homologue of ZBP1, the insulin-like growth factor-2 (IGF2) binding protein, *IGF2BP1-3* which has been characterized as an m^6^A reader and has a regulatory effect on *MYC* expression. However, in this earlier study, IGF2BP1-3 binding was tested not with the actin zipcode but with tandem repeats of GGACU which had multiple m^6^A modifications (Huang et al. 2018).

In addition, we found that replacing As with m^6^A at other sites within the 28nt core zipcode sequence could also influence ZBP1KH3-KH4 binding. Out of three different positions, one (A16), did not change the ZBP1 binding compared to the unmodified zipcode. The two remaining positions had negative effects leading to a nearly complete loss of binding when m^6^A replaced the A in the **<u>A</u>**CACCCC (position 22, E oligonucleotide) motif. This motif is also an essential component for the ZBP1KH3-KH4 binding, as changing this adenosine to a guanosine (position 22) was previously shown to disrupt the ZBP1KH3-KH4-RNA complex formation (Chao et al. 2010). In a recent study, this sequence region was also shown to bind specifically to a KH3 domain, whilst the KH4 domain was responsible for binding to the GA in the GG**<u>A</u>**CU motif (Nicastro et al. 2017). KH3 preferentially binds to AC rich regions via the C in position 23. The A following the C, in position 24 can be replaced with a C without significantly changing the Kd value. The effects of replacing A in position 22, was not examined in this study (Nicastro et al. 2017). The two previous studies (Chao et al. 2010, Huang et al. 2018) and our gel shift assays demonstrate that the zipcode domains **<u>A</u>**CACCCC and GG**<u>A</u>**CU are important for ZBP1 binding. The **<u>m^6^A</u>**CACCCC decreases, while GG**<u>m^6^A</u>**CU increases, ZBP1 binding *in vitro*. Thus, we hypothesize that the presence of an **<u>m^6^A</u>**CACCCC motif may counteract the effect of GG**<u>m^6^AC</u>**U *in vivo*, this could have significant consequences for the regulation of β-actin expression, and localisation. However, we did not detect any m^6^A at the **<u>A</u>**CACCCC site, and found that 14% of As were methylated at the GG**<u>A</u>**CU position in chicken embryos. We also showed that the m^6^A is present at the GG**<u>A</u>**CU position in embryonal fibroblast cells at a higher stoichiometry (20%).

The zipcode controlled localisation of *ACTB* mRNA determines cell polarity and mobility in chicken embryonal fibroblast cell as well as other cell types (Shestakova et al. 2001). This process is facilitated by ZBP1 binding. Our results suggest that ZBP1 binding to the core zipcode sequence can be altered by differential m^6^A deposition. This highlights the importance of accurate m^6^A deposition by the writer complex and also of the maintenance of a dynamic equilibrium between the m^6^A writing and erasing *in vivo*. This also underlines the importance of knowing the precise topology of the m^6^A molecule at the single transcript level and emphasizes the need for utilising RNA oligonucleotides with modifications at defined positions when carrying out RNA-protein binding assays, rather than RNA substrates generated by transcription in the presence of the modified nucleoside triphosphate, allowing a multiple but untargeted incorporation of m^6^A.

### An improved method for m^6^A site verification and quantification

The presence of m^6^A is essential for the control and fine tuning of multiple cell differentiation and developmental processes in all eukaryotes where it has been studied. In most eukaryotes multiple m^6^A sites are frequently observed at a single transcript level (Meyer et al. 2012) and the depletion of m^6^A can give rise to pleiotropic effects. Thus, m^6^A removal may result in diverse, or no effect, on the function of a single mRNA molecule depending on the position from where the m^6^A was removed. The topology of m^6^A at single nucleotide resolution, and its stoichiometry in a transcript population, are therefore fundamental to our understanding of the m^6^A functional consequences.

Unequivocal single nucleotide resolution is not possible from MeRIPSeq data as the peaks are broad and the peak summits do not always fall over the m^6^A position. Indeed, many m^6^A calling pipelines look for the nearest RRACH under or close to the peak summit (Schwartz et al. 2014). It was previously claimed that by increased fragmentation of the RNA and more refined bioinformatics approaches, a near single nucleotide resolution may be feasible. (Schwartz et al. 2014). Increased resolution by simply refining bioinformatics is not possible without improved accuracy and specificity of m^6^A detection chemistry. Thus, development of antibody independent biochemical verification methods are essential. Recent studies utilizing Oxford Nanopore sequencing show promise in detecting RNA modifications (Liu et al. 2019, Parker et al. 2020). However, such approaches are still in development, therefore these methods would benefit from an independent and unbiased approach to enable authentication of specific m^6^A positions by direct biochemical methods.

This study addresses the need for a biochemical method that detects and precisely and unambiguously identifies m^6^A in any RNA molecule with very high confidence. The antibody independent SCARLET (Liu et al. 2013) can detect m^6^A site-specifically at the transcript level, however this method has not been widely adopted for routine laboratory use. Despite its elegance, SCARLET requires lengthy preparatory steps and gel purification that substantially decreases the yield of final product, and thus necessitates increased amounts of starting material that are not always feasible. Furthermore, SCARLET uses a targeted RNaseH cleavage (Zhao and Yu, 2004) at the m^6^A site, leaving an RNA fragment with an exposed 5′pA/pm^6^A end. The following steps require removal of this 5′ phosphate and addition of a labelled 5’ phosphate. This is followed by creating DNA RNA chimera to enable gel purification of the target RNA fragments using T4 DNA ligase and a splint DNA specific to the ends of the labelled m^6^A RNA and DNA molecules. However, the activity of T4 DNA ligase is not blocked by the presence of gaps between the nucleic acid ends and the enzyme is able to carry out the ligation process (Lohman et al. 2014). Thus ends from misclevage by RNaseH are likely to be labelled and gel purified. One of these ends could be the C following the A/m^6^A. This labeled C can be misinterpreted as m^6^A on the one-dimensional TLC used in the SCARLET method, as pC would run at a very similar Rf value to the pm^6^A under the applied conditions (buffer used for one dimensional TLCs). The RedBaron method avoids these artefacts by using SplintR^®^Ligase that can only ligate if there is no gap in the double stranded region between the 3′ and 5′ ends of contributing molecules. The RedBaron does not need gel purification and requires relatively low input amounts of RNA. In addition, our method uses two dimensional TLC, thus giving unequivocal resolution of nucleotide spots and avoiding potential miscalling of nucleotides with similar Rf values in the first dimension buffer. The biochemical steps can be performed in a day, which is a significant improvement over more complex methods. RedBaron is also reproducible, accurate and a low-input method for m^6^A site verification. Furthermore, RedBaron, unlike SELECT, does not need an accurate quantification of the input RNA, or calibration curves and concentration of the target transcript for determining the m^6^A/A ratios.

In the field of m^6^A epitranscriptomics, much attention has been given to the conserved YT521-B homology (YTH) domain-containing proteins that preferentially bind m^6^A and act as “readers” of methylated transcripts. However, the work reported here and elsewhere shows that other proteins also act as m^6^A readers via their enhanced binding when the modification is present. Importantly, it also highlights that RNA binding can be abolished by the presence of m^6^A and furthermore, enhanced or inhibited RNA binding of a given protein can be m^6^A context dependent.

Plasticity of the actin cytoskeleton is important for many cell and developmental processes, including stem cell differentiation; and defective β-actin localization can promote cancer metastasis (Shestakova et al. 1999). Altered mRNA methylation has been associated with faulty cell differentiation and with cancer progression. We would like to suggest that the potential involvement of aberrant β-actin localization should be considered in some of these cases.

## Materials and Methods

### Cells and tissues

Chicken embryonal fibroblast cells (DF1; a kind gift from Dr Dylan Sweetman) were expanded in standard Dulbecco’s Modified Eagle Medium (DMEM; Merck Life Science UK Limited, Gillingham, Dorset, UK) supplemented with 10% fetal bovine serum (FBS; Merck) and 5% penicillin/streptomycin (Merck). Cells were split at 80% confluence using trypsin-ethylenediaminetetraacetic acid (trypsin EDTA; Merck), spun at 200 x g for 5 minutes to pellet and snap frozen in liquid nitrogen. Cell pellets were stored at −80°C. Fertile chicken eggs (*Gallus gallus*; Henry Stewart, UK) were incubated and the chicken embryos were collected after 5.5 days at Hamburger and Hamilton stage 27 (HH 27) (Hamburger and Hamilton, 1951). The samples were snap frozen in liquid nitrogen and stored at –80°C. The work was performed within national (UK Home Office) and institutional ethical regulations with permission from the School of Veterinary Medicine and Science ethics committee (ethics number 2320 180612). The HeLa-S3-Cells total RNA was purchased from Agilent Technologies.

### RNA purification

Total RNA was prepared from cells and tissues using Trizol reagent (Invitrogen). The poly(A) RNA was prepared using two rounds of oligo(dT) magnetic beads purification (New England Biolabs).

### RedBaron method

The ssDNA oligonucleotide (Table S3) (15 pmol) was mixed with ATP [γ-^32^P] (16 pmol, 48 μCi), and T4 Poly Nucleotide Kinase (10 U) (New England Biolabs) in a total volume of 30 μL PNK buffer A (1x). The solution was incubated at 37 °C for 1 hr, followed by 75 °C for 5 mins. The radiolabeled oligonucleotide was purified using a QIAquick Nucleotide Removal Kit and eluted in 100 μL H_2_O. An 18-22 nts chimeric nucleotide was designed to give an appropriate melting temperature, and in the center containing 4 DNA nucleotides covering the RNase H cut site.

Poly(A)+ RNA (100 ng) was mixed with the chimeric oligonucleotide (Table S3) (1 pmol) in a volume of 27.5 µL Tris-HCl (30 mM, pH 7.5). The nucleic acid was annealed by incubating at 95°C for 1 min, followed by room temperature for 5 min. 1.5 µL PNK buffer (10x) (New England Biolabs) was added, and the solution was incubated at 44 °C for 5 min. 1 µL RNase H enzyme (5 U) (New England Biolabs) was added and the solution was incubated at 44°C for 1 hr. The solution was mixed with TRIzol Reagent (Invitrogen) (500 µL) and incubated at room temperature for 3 min. The solution was then mixed with chloroform (500 µL) and incubated at room temperature for 2 min. The solution was centrifuged at 13,000 x g for 15 min and the upper aqueous phase was mixed with an equal volume of ethanol. The RNA was purified using an RNA Clean & Concentrator-5 Kit (Zymo Research) and eluted into 10 µL H_2_O.

The RNase H treated RNA (10 µL) was mixed with the 5′ ^32^P radiolabeled ssDNA oligonucleotide (1.5 pmol, 4.5 µCi), and the splint oligonucleotide (Table S3) (1 pmol) in a total volume of 26 µL Tris-HCl (30 mM, pH 7.5). The nucleic acid was annealed by incubating the solution at 75°C for 3 min followed by room temperature for 5 min. 3 µL SplintR® Ligase buffer (10x) was added and the solution was incubated at 37°C for 5 min. 1 µL SplintR® Ligase (25 U) (New England Biolabs) was added and the solution was incubated at 37°C for 1 hr followed by 75°C for 5 min, and 5 min on ice. 2 µL FastAP Thermosensitive Alkaline Phosphatase (2 U) (Thermo Scientific) was added and the solution was incubated at 37°C for 20 min, followed by 75°C for 5 min. The nucleic acid was purified using a RNA Clean & Concentrator-5 (Zymo Research) and eluted in 7µL H_2_O.

7 μL of the nucleic acid was mixed with 1 μL BSA (10 x), 1 μL Micrococcal Nuclease buffer (10x), and 1 μL Micrococcal Nuclease (2000 U) (New England Biolabs). The solution was incubated at 37°C for at least 3-4 hrs. 1 µL of the solution was spotted onto a TLC Cellulose glass plate (20×20 cm). The TLC was resolved in two dimensions and imaged and quantified using a FX Phospho imager (Bio-Rad Laboratories) in combination with the QuantityOne 4.6.8 software. For the synthesis of 5′ ^32^P radiolabeled mononucleotide reference molecules, the in-house oligonucleotide synthesis and the synthesis of 2′-OTBS-Bz-m6A-CE phosphoramidite see Appendix Supplementary Materials and Methods.

### MeRIP-Seq

Total RNA from 3 replicates of chicken embryos stage HH27 was isolated as previously described. This was followed by one round poly(A) purification using oligo d(T) magnetic beads (New England Biolabs). 1.5-2 µg of mRNA was fragmented to 100-150 nts using RNA Fragmentation Reagent (Thermo Fisher Scientific) followed by overnight ethanol precipitation. After centrifugation and washing, the pellets were resuspended in 10 µL H_2_O. 9 μL of the solution was used for the IP, and 1 µL for preparing the input libraries. The fragmented RNA was mixed with Protein G Magnetic bead prebound monoclonal anti-m^6^A antibody (1 µL) from the EpiMark *N*6-Methyladenosine Enrichment Kit (New England Biolabs), resuspended in 300 µl EpiMark IP buffer supplemented with murine RNase inhibitor (New England Biolabs). All following steps were as described by the manufacturer. After the last wash we carried out an extra washing step using H_2_O. We omitted the final elution step as we carried out the cDNA synthesis on the magnetic beads using ScriptSeq v2 RNA-Seq Library Preparation Kit (Illumina). The libraries were size selected using BluePippin DNA size selection system (Sage Science), and quality checked on Agilent High Sensitivity DNA Chips (Agilent Technologies). The pooled libraries were sequenced on Nextseq 500 (Illumina) DeepSeq at The University of Nottingham.

### Sequencing analysis

Contaminating adapter sequences and low quality reads (phred scores <30) were removed using TrimGalore(v0.4.4). The processed fastq reads were aligned to the Ensembl annotated Chicken GRCg6a reference genome using STAR(v2.5.0), the resultant bam files were indexed using Samtools (v1.10, PMID:19505943) and m^6^A enriched regions identified in m^6^A immunoprecipitated samples over inputs, using m6AViewer (v1.6.1) (PMID:<u>28724534</u>) (Antanaviciute et al. 2017). Bedtools (v2.27.1, PMID:20110278) was used to extend peaks by 100bp upstream and downstream. Only those significant peaks represented in at least two replicates, and 4-fold enriched were taken forward for further analysis from the peak matrix dataset generated in the RNAmod software (http://61.147.117.195/RNAmod) with default settings (Liu and Gregory, 2019). Using the Peak matrix dataset created by the RNAmod software, for single replicates a KEGG pathway enrichment analysis was carried out by this online tool. In addition DAVID Bioinformatics Resources (Huang et al. 2009) (https://david.ncifcrf.gov/) was used for the KEGG pathway analysis of the 3’UTR methylated chicken transcripts conserved between mouse and chicken. For finding conserved methylated transcripts between chicken and mouse we downloaded the complete gene list of all vertebrate homologues from MGI (http://www.informatics.jax.org/homology.shtml) and the peak files for mouse embryoid bodies (Geula et al. 2015) from REPIC (https://repicmod.uchicago.edu/repic) (Liu et al. 2020).

### Gel shift assay

To assess the binding of the modified and unmodified zipcode RNA oligonucleotides to the recombinant ZBP1KH3-KH4 protein we carried out gel shift assays. The RNA oligonucleotides (Table S3) were end labelled using ATP [γ-^32^P] and T4 polynucleotide kinase, followed by purification using a QIAquick Nucleotide Removal Kit (QIAGEN) and eluted with H_2_O. The recombinant ZBP1KH3-KH4 (16 nM) and the RNA oligonucleotides (Table S3) (2.5 nM) were incubated for 3 h at the same conditions described by Chao et al. (2010). The protein-RNA complexes were resolved on 5% TBE polyacrylamide precast gels (Bio-Rad Laboratories), and for imaging purposes were transferred onto Hybond-N membranes (GE Healthcare) followed by an exposure to phosphor screen (FUJI). The scanned images (FX scanner Bio-Rad Laboratories) were quantified using the QuantityOne 4.6.8 software (Bio-Rad Laboratories).

### Protein expression

The PCR product of the truncated ZBP1KH3-Kh4, 404–561 with a C terminal His tag added was cloned into the pMAL-c5x vector (New England Biolabs), and were transformed in *E. coli* (DH5α). The recombinant protein was induced with 2% ethanol and 0.4mM IPTG and grown for 20 hours at 18°C. The cells were pelleted by centrifugation and resuspended in Ni column equilibration buffer (20mM Na_3_PO_4_; 300mM NaCl, 10mM Imidazole at pH 7.4) and lysed by sonication. ZBP1KH3-KH4 was purified using the gravity flow column with HisPur Ni-NTA resin (Fisher Scientific).

## Data availability

All data are accessible from NCBI under the GEO accession number: GSE185078

## Acknowledgments

Work in the laboratories of RGF, CJH, HMK, CSR, and NPM was supported by the Biotechnology and Biological Sciences Research Council Doctoral Training Program (BB/I024291/1) to FB and EB. NPM supported by Prostate Cancer Foundation-John Black Charitable Foundation Challenge Award (20CHAL04) and RGF by BBSRC (Grant BB/K013637/1). Work in CSR’s laboratory was supported by a Faculty of Medicine and School of Veterinary Medicine and Science, University of Nottingham strategic grant. NA is a Nottingham Research Fellow funded by the University of Nottingham.

## References

Almuzzaini B, Sarshad AA, Rahmanto AS, Hansson ML, Von Euler A, Sangfelt O, Visa N, Farrants AK, Percipalle P. 2016. In β-actin knockouts, epigenetic reprogramming and rDNA transcription inactivation lead to growth and proliferation defects. FASEB J 30: 2860–2873. doi:10.1096/fj.201600280R

Antanaviciute A, Baquero-Perez B, Watson CM, Harrison SM, Lascelles C, Crinnion L, Markham AF, Bonthron DT, Whitehouse A, Carr IM. 2017. m6aViewer: software for the detection, analysis, and visualization of N^6^-methyladenosine peaks from m^6^A-seq/ME-RIP sequencing data. RNA 23: 1493–1501. doi:10.1261/rna.058206.116

Aschenbrenner J, Werner S, Marchand V, Adam M, Motorin Y, Helm M, Marx A. 2018. Engineering of a DNA polymerase for direct m^6^A sequencing. Angew Chem Int Ed 57: 417–421. doi:10.1002/anie.201710209

Chao JA, Patskovsky Y, Patel V, Levy M, Almo SC, Singer RH. 2010. ZBP1 recognition of β-actin zipcode induces RNA looping. Genes Dev 24: 148–158. doi:10.1101/gad.1862910

Chen M, Wei L, Law CT, Tsang FH, Shen J, Cheng CL, Tsang LH, Ho DW, Chiu DK, Lee JM, et al. 2018. RNA N6-methyladenosine methyltransferase-like 3 promotes liver cancer progression through YTHDF2-dependent posttranscriptional silencing of SOCS2. Hepatology 67: 2254– 2270. doi:10.1002/hep.29683

Dominissini D, Moshitch-Moshkovitz S, Schwartz S, Salmon-Divon M, Ungar L, Osenberg S, Cesarkas K, Jacob-Hirsch J, Amariglio N, Kupiec M, et al. 2012. Topology of the human and mouse m^6^A RNA methylomes revealed by m^6^A-seq. Nature 485: 201–206. doi:10.1038/nature11112

Du H, Zhao Y, He J, Zhang Y, Xi H, Liu M, Ma J, Wu L. 2016. YTHDF2 destabilizes m^6^A-containing RNA through direct recruitment of the CCR4–NOT deadenylase complex. Nat Commun 7: 12626. doi:10.1038/ncomms12626

Geula S, Moshitch-Moshkovitz S, Dominissini D, Mansour AA, Kol N, Salmon-Divon M, Hershkovitz V, Peer E, Mor N, Manor YS, et al. 2015. m^6^A mRNA methylation facilitates resolution of naïve pluripotency toward differentiation. Science 347: 1002–1006. doi:10.1126/science.1261417

Hamburger V, Hamilton HL. 1951. A series of normal stages in the development of the chick embryo. Dev Dyn 195: 231–272.

Harcourt EM, Ehrenschwender T, Batista PJ, Chang HY, Kool ET. 2013. Identification of a selective polymerase enables detection of N^6^-methyladenosine in RNA. J Am Chem Soc 135: 19079–19082. doi:10.1021/ja4105792

Helm M, Lyko F, Motorin Y. 2019. Limited antibody specificity compromises epitranscriptomic analyses. Nat Commun 10: 5669. doi:10.1038/s41467-019-13684-3

Hong T, Yuan Y, Chen Z, Xi K, Wang T, Xie Y, He Z, Su H, Zhou Y, Tan ZJ, et al. 2018. Precise antibody-independent m6A identification via 4SedTTP-involved and FTO-assisted strategy at single-nucleotide resolution. J Am Chem Soc 140: 5886–5889. doi:10.1021/jacs.7b13633

Huang H, Weng H, Sun W, Qin X, Shi H, Wu H, Zhao BS, Mesquita A, Liu C, Yuan CL. 2018. Recognition of RNA N^6^-methyladenosine by IGF2BP proteins enhances mRNA stability and translation. Nat Cell Biol 20: 285–295. doi:10.1038/s41556-018-0045-z

Huang da W, Sherman BT, Lempicki RA. 2009. Systematic and integrative analysis of large gene lists using DAVID bioinformatics resources. Nature Protoc 4: 44–57. doi:10.1038/nprot.2008.211

Imanishi M, Tsuji S, Suda A, Futaki S. 2017. Detection of N^6^-methyladenosine based on the methyl-sensitivity of Maz F RNA endonuclease. Chem Commun 50: 12930–12933. doi:10.1039/C7CC07699A

Ke S, Alemu EA, Mertens C, Gantman EC, Fak JJ, Mele A, Haripal B, Zucker-Scharff I, Moore MJ, Park CY, Vågbø CB, et al. 2015. A majority of m^6^A residues are in the last exons, allowing the potential for 3′ UTR regulation. Genes Dev 29: 2037–2053. doi:10.1101/gad.269415.115

Lawrence JB, Singer RH. 1986. Intracellular localization of messenger RNAs for cytoskeletal proteins. Cell 45: 407–415. doi:10.1016/0092-8674(86)90326-0

Lehtimäki J, Hakala M, Lappalainen P. 2017. Actin filament structures in migrating cells. Handb Exp Pharmacol 235: 123–152. doi:10.1007/164_2016_28

Lesbirel S, Viphakone N, Parker M, Parker J, Heath C, Sudbery I, Wilson SA. 2018. The m^6^A-methylase complex recruits TREX and regulates mRNA export. Sci Rep 8: 13827. doi:10.1038/s41598-018-32310-8

Linder B, Grozhik AV, Olarerin-George AO, Meydan C, Mason CE, Jaffrey SR. 2015. Single-nucleotide-resolution mapping of m6A and m6Am throughout the transcriptome. Nat Methods 12: 767–772. doi:10.1038/nmeth.3453

Liu H, Begik O, Lucas MC, Ramirez JM, Mason CE, Wiener D, Schwartz S, Mattick JS, Smith MA, Novoa EM. 2019. Accurate detection of m^6^A RNA modifications in native RNA sequences. Nat Commun 10: 4079. doi:10.1038/s41467-019-11713-9

Liu J, Li K, Cai J, Zhang M, Zhang X, Xiong X, Meng H, Xu X, Huang Z, Peng J,et al. 2020. Landscape and Regulation of m6A and m6Am Methylome across Human and Mouse Tissues. Mol Cell. 77: 426-440.e6. doi: 10.1016/j.molcel.2019.09.032.

Liu N, Parisien M, Dai Q, Zheng G, He C, Pan T. 2013. Probing N^6^-methyladenosine RNA modification status at single nucleotide resolution in mRNA and long noncoding RNA. RNA 19: 1848–1856. doi:10.1261/rna.041178.113

Liu S, Zhu A, He C, Chen M. 2020. REPIC: a database for exploring the N^6^-methyladenosine methylome. Genome Biol 21: 100. doi:10.1186/s13059-020-02012-4

Liu Q, Gregory RI. 2019. RNAmod: an integrated system for the annotation of mRNA modifications. Nucleic Acids Res 47: W548–W555. doi:10.1093/nar/gkz479

Lohman GJ, Zhang Y, Zhelkovsky AM, Cantor EJ, Evans TC Jr. 2014. Efficient DNA ligation in DNA–RNA hybrid helices by Chlorella virus DNA ligase. Nucleic Acids Res 42: 1831–1844. doi:10.1093/nar/gkt1032

Lu Y, Li S, Zhu S, Gong Y, Shi J, Xu L. 2017. Methylated DNA/RNA in body fluids as biomarkers for lung cancer. Biol Proced Online 19: 2. doi:10.1186/s12575-017-0051-8

Luxenburg C, Geiger B. 2017. Multiscale view of cytoskeletal mechanoregulation of cell and tissue polarity. Handb Exp Pharmacol 235: 263–284. doi: 10.1007/164_2016_34

Meyer KD, Saletore Y, Zumbo P, Elemento O, Mason CE, Jaffrey SR. 2012. Comprehensive analysis of mRNA methylation reveals enrichment in 3′ UTRs and near stop codons. Cell 149: 1635–1646. doi:10.1016/j.cell.2012.05.003

Meyer KD, Patil DP, Zhou J, Zinoviev A, Skabkin MA, Elemento O, Pestova TV, Qian SB, Jaffrey SR. 2015. 5′ UTR m^6^A promotes cap-independent translation. Cell 163: 999–1010. doi:10.1016/j.cell.2015.10.012

Nicastro G, Candel AM, Uhl M, Oregioni A, Hollingworth D, Backofen R, Martin SR, Ramos A. 2017. Mechanism of β-actin mRNA recognition by ZBP1. Cell Rep 18: 1187–1199. doi:10.1016/j.celrep.2016.12.091

Parker MT, Knop K, Sherwood AV, Schurch NJ, Mackinnon K, Gould PD, Hall AJ, Barton GJ, Simpson GG. 2020. Nanopore direct RNA sequencing maps the complexity of Arabidopsis mRNA processing and m^6^A modification. eLife 9: e49658. doi:10.7554/eLife.49658

Roundtree IA, Luo GZ, Zhang Z, Wang X, Zhou T, Cui Y, Sha J, Huang X, Guerrero L, Xie P, et al. 2017. YTHDC1 mediates nuclear export of N^6^-methyladenosine methylated mRNAs. eLife 6: e31311. doi:10.7554/eLife.31311

Schwartz S, Mumbach MR, Jovanovic M, Wang T, Maciag K, Bushkin GG, Mertins P, Ter-Ovanesyan D, Habib N, Cacchiarelli D, et al. 2014. Perturbation of m6A writers reveals two distinct classes of mRNA methylation at internal and 5′ sites. Cell Rep 8: 284–296. doi:10.1016/j.celrep.2014.05.048

Shestakova EA, Wyckoff J, Jones J, Singer RH, Condeelis J. 1999. Correlation of beta-actin messenger RNA localization with metastatic potential in rat adenocarcinoma cell lines. Cancer Res. 59: 1202–1205.

Shestakova EA, Singer RH, Condeelis J. 2001. The physiological significance of β-actin mRNA localization in determining cell polarity and directional motility. Proc Natl Acad Sci USA 98: 7045–7050. doi:10.1073/pnas.121146098

Vedula P, Kashina A. 2018. The makings of the ‘actin code’: regulation of actin’s biological function at the amino acid and nucleotide level. J Cell Sci 131: 9. doi:10.1242/jcs.215509

Viita T, Vartiainen MK. 2017. From cytoskeleton to gene expression: actin in the nucleus. Handb Exp Pharmacol 235: 311–329. doi:10.1007/164_2016_27

Vilfan ID, Tsai Y-C, Clark TA, Wegener J, Dai Q, Yi C, Pan T, Turner SW, Korlach J. 2013. Analysis of RNA base modification and structural rearrangement by single-molecule real-time detection of reverse transcription. J Nanobiotechnol 11: 8. doi:10.1186/1477-3155-11-8

Wang S, Wang J, Zhang X, Fu B, Song Y, Ma P, Gu K, Zhou X, Zhang X, Tian T, et al. 2016. N^6^-methyladenine hinders RNA- and DNA-directed DNA synthesis: application in human rRNA methylation analysis of clinical specimens. Chem Sci 7: 1440–1446. doi:10.1039/C5SC02902C

Wang X, Lu Z, Gomez A, Hon GC, Yue Y, Han D, Fu Y, Parisien M, Dai Q, Jia G, et al. 2014. N^6^-methyladenosine-dependent regulation of messenger RNA stability. Nature 505: 117–120. doi:10.1038/nature12730

Wang X, Zhao BS, Roundtree IA, Lu Z, Han D, Ma H, Weng X, Chen K, Shi H, He C. 2015. N^6^-methyladenosine modulates messenger RNA translation efficiency. Cell 161: 1388–1399. doi:10.1016/j.cell.2015.05.014

Xiao Y, Wang Y, Tang Q, Wei L, Zhang X, Jia G. 2018. An elongation- and ligation-based qPCR amplification method for the radiolabeling-free detection of locus-specific N^6^-methyladenosine modification. Angew Chem Int Ed 57: 15995–16000. doi:10.1002/anie.201807942

Xiao W, Adhikari S, Dahal U, Chen YS, Hao YJ, Sun BF, Sun HY, Li A, Ping XL, Lai WY, Wang X, et al. 2016. Nuclear m^6^A reader YTHDC1 regulates mRNA splicing. Mol Cell 61: 507– 519. doi:10.1016/j.molcel.2016.01.012

Zhang C, Samanta D, Lu H, Bullen JW, Zhang H, Chen I, He X, Semenza GL. 2016. Hypoxia induces the breast cancer stem cell phenotype by HIF-dependent and ALKBH5-mediated m^6^A-demethylation of NANOG mRNA. Proc Natl Acad Sci USA 113: 2047–2056. doi:10.1073/pnas.1602883113

Zhang HL, Singer RH, Bassell GJ. 1999. Neurotrophin regulation of β-Actin mRNA and protein localization within growth cones. J Cell Biol 147: 59–70. doi:10.1083/jcb.147.1.59

Zhang Z, Chen LQ, Zhao YL, Yang CG, Roundtree IA, Zhang Z, Ren J, Xie W, He C, Luo GZ. 2019. Single-base mapping of m^6^A by an antibody-independent method. Sci Adv 5: 10. doi:10.1126/sciadv.aax0250

Zhao X, Yu YT. 2004. Detection and quantitation of RNA base modifications. RNA 10: 996–1002. doi:10.1261/rna.7110804

Zhong S, Li H, Bodi Z, Button J, Vespa L, Herzog M, Fray RG. 2008. MTA is an Arabidopsis messenger RNA adenosine methylase and interacts with a homolog of a sex-specific splicing factor. Plant Cell 20: 1278–1288. doi:10.1105/tpc.108.058883

Zhou J, Wan J, Gao X, Zhan X, Jaffrey SR, Qian SB. 2015. Dynamic m^6^A mRNA methylation directs translational control of heat shock response. Nature 526: 591–594. doi:10.1038/nature15377

